# In Vivo Bioincubation Promotes Maturation of Human iPSC-Derived Cardiomyocytes in Neonatal Rat and Pig Hearts

**DOI:** 10.64898/2026.07.21.739858

**Authors:** Hanwen Wang, Peter Andersen, Taka Inoue, Narutoshi Hibino, Dong Ik Lee, Chulan Kwon

## Abstract

Human induced pluripotent stem cell-derived cardiomyocytes (hiPSC-CMs) hold great promise for cardiac regenerative medicine and disease modeling. However, hiPSC-CMs generated through conventional *in vitro* differentiation exhibit immature, fetal-like phenotypes. While *in vivo* bioincubation in neonatal rodent hearts promotes hiPSC-CM maturation toward adult-like phenotypes, studies in large animal models remain limited, particularly with detailed morphological characterization. In this study, we investigated bioincubation of fluorescently labeled hiPSC-CMs in both neonatal rat and pig hearts. Human iPSCs were differentiated into cardiomyocytes expressing GFP or RFP reporters and subsequently injected intramyocardially into neonatal rats (GFP-labeled) and pigs (RFP-labeled). After 4–8 weeks of bioincubation, fluorescent hiPSC-CMs were isolated using large-particle fluorescence-activated cell sorting (COPAS), which preserves cellular morphology of adult-like cardiomyocytes. Immunostaining for cardiac troponin T revealed well-organized sarcomeric structures in multinucleated hiPSC-CMs. Bioincubated hiPSC-CMs displayed rod-shaped morphology with binucleation, characteristic features of mature adult cardiomyocytes. Quantitative analysis demonstrated that bioincubated hiPSC-CMs from rat hearts exhibited sarcomere length and cell circularity comparable to native rat adult cardiomyocytes, though with higher intra-cellular variability in sarcomere organization. Histological examination confirmed successful engraftment of RFP-positive hiPSC-CMs within pig myocardium, with engrafted cells also displaying mature adult-like features. These findings provide critical proof-of-concept data for bioincubation in large animal models and support further investigation for disease modeling, drug screening, and regenerative cell therapies.

**SIGNIFICANCE STATEMENT:** Human induced pluripotent stem cell-derived cardiomyocytes (hiPSC-CMs) offer tremendous potential for cardiac disease modeling and regenerative therapies, but their clinical application is limited by their immature characteristics. Here we show that *in vivo* bioincubation in neonatal rat hearts enables hiPSC-CMs to achieve structural maturity, exhibiting features of adult cardiomyocytes, including organized sarcomeres, rod-shaped morphology, and multinucleation. We further provided proof-of-concept evidence for engraftment in neonatal pig hearts for maturation, supporting feasibility in large animal models. The use of large-particle cell sorting enables recovery of intact, adult-sized cardiomyocytes for subsequent analysis. These findings establish a practical and scalable platform for generating structurally mature human cardiomyocytes through *in vivo* bioincubation.

## INTRODUCTION

Human induced pluripotent stem cell-derived cardiomyocytes (hiPSC-CMs) have emerged as a transformative platform for cardiac disease modeling, drug discovery, and regenerative medicine [1, 2]. These cells offer valuable opportunities to study patient-specific cardiac pathophysiology and develop personalized therapeutic strategies. However, hiPSC-CMs generated through current differentiation protocols remain developmentally immature, exhibiting fetal-like characteristics that fundamentally differ from adult cardiomyocytes [1–3]. This immaturity manifests as disorganized myofibrils, reduced anisotropy, absence of transverse tubules, underdeveloped excitation-contraction coupling, and electrophysiological properties distinct from adult myocardium. During the critical perinatal window, native cardiomyocytes undergo coordinated structural, functional, and metabolic transitions that establish the adult phenotype [4]. hiPSC-CMs fail to complete these developmental programs, remaining arrested at an embryonic-like state that limits their utility for modeling adult-onset cardiomyopathies [4–6].

Various *in vitro* approaches have been developed to enhance hiPSC-CM maturation, including prolonged culture, electrical stimulation, mechanical loading, and three-dimensional culture systems [8–10]. These approaches have yielded incremental improvements but still fall short of recapitulating the complex maturation cues present in the developing heart. Functional, metabolic, and transcriptional aspects of cardiomyocyte maturation can proceed independently, suggesting that comprehensive maturation requires integrated signals that may only exist *in vivo* [11]. Kadota et al. showed that hiPSC-CMs transplanted into adult rat hearts developed organized sarcomeres and increased cell size over 3 months, with cell area increasing from approximately 97 *µ*m^2^ to 247 *µ*m^2^ [12]. Cho et al. demonstrated that neonatal transplantation in rats confers maturation of PSC-CMs to a degree conducive to modeling inherited cardiomyopathies [13].

Despite progress in rodent models, bioincubation in large animal models remains unexplored. Compared to small rodents, large animals such as pigs more closely approximate human cardiac physiology, making them essential for translational validation. Recovery of matured hiPSC-CMs from host tissue also presents technical barriers, as conventional cell sorting damages large, fragile adult cardiomyocytes. In this study, we demonstrated bioincubation of fluorescently labeled hiPSC-CMs in both neonatal rat and pig hearts and employed COPAS large-particle sorting to isolate intact adult-sized cardiomyocytes.

## MATERIALS AND METHODS

### Human iPSC Culture and Cardiac Differentiation

Human iPSCs (WTC11) were maintained on Matrigel-coated plates in mTeSR Plus medium under standard culture conditions (37 °C, 5% CO_2_). For fluorescent labeling, hiPSCs were transduced with lentiviral vectors encoding constitutive CAG promoter-driven GFP (for rat studies) or RFP (for pig studies). Cardiac differentiation was performed using temporal modulation of Wnt signaling with small molecules. Briefly, hiPSCs at approximately 85% confluence were treated with CHIR99021 to induce mesoderm formation, followed by IWR1 to promote cardiac specification. Cardiomyocyte purity was assessed by flow cytometry for cardiac troponin T (cTnT), and batches with >80% cTnT-positive cells were used for transplantation.

### In Vivo Bioincubation

All procedures were approved by the Johns Hopkins University Institutional Animal Care and Use Committee. Neonatal immunodeficient rats at postnatal day 2–3 received intramyocardial injection of 1 million GFP-labeled hiPSC-CMs into the left ventricular free wall as reported [6, 7]. Animals were maintained for 4 weeks before heart harvest. Piglets received intramyocardial injection of 1 million RFP-labeled hiPSC-CMs into the left ventricular myocardium and were monitored for 8 weeks before histological analysis.

### Cardiomyocyte Isolation and COPAS Sorting

Following bioincubation, rat hearts were excised and enzymatically digested to generate single-cell suspensions. The cell suspension was subjected to large-particle fluorescence-activated cell sorting using a COPAS VISION system (Union Biometrica). GFP-positive hiPSC-CMs and GFP-negative native rat cardiomyocytes were collected separately for downstream analysis.

### Immunofluorescence and Morphometric Analysis

Sorted cardiomyocytes were plated on laminin-coated coverslips, fixed with 4% paraformaldehyde, and immunostained for cTnT with DAPI nuclear counterstain. Sarcomere length was quantified using Haralick correlation analysis of cTnT staining patterns. Cell circularity (4*π ×* area/perimeter^2^) and cell area were measured using ImageJ software.

### Statistical Analysis

Data are presented as mean *±* SEM. Comparisons between groups were performed using unpaired Student’s *t*-test or Mann-Whitney U test as appropriate. *P* < 0.05 was considered statistically significant.

## RESULTS

To enable tracking and recovery of hiPSC-CMs following *in vivo* bioincubation, we generated stable reporter lines by transducing human iPSCs with constitutive CAG-driven GFP or RFP (Fig. 1a). Directed cardiac differentiation produced spontaneously contracting cardiomyocyte populations.

**Figure 1.**
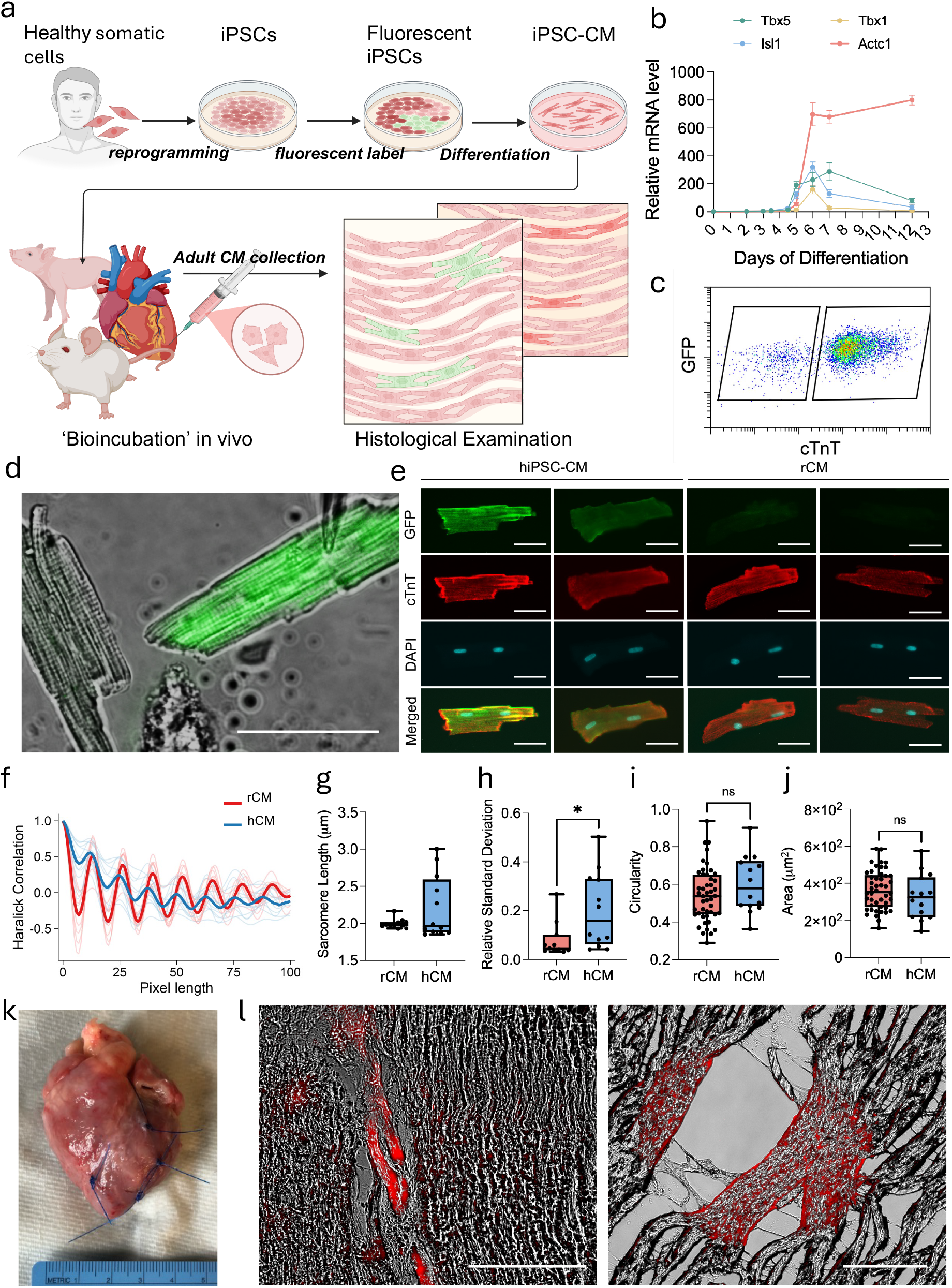
Bioincubation of human iPSC-derived cardiomyocytes promotes maturation *in vivo*. **(a)** Schematic of the bioincubation workflow. Human iPSCs were reprogrammed from healthy somatic cells, transduced with constitutive fluorescent reporters (CAG-GFP or CAG-RFP), and differentiated into cardiomyocytes (iPSC-CMs). Fluorescent iPSC-CMs were delivered via intramy-ocardial injection into neonatal rat or pig hearts for *in vivo* bioincubation. Following bioincubation, hearts were harvested for enzymatic digestion and COPAS sorting (rat) or histological examination (pig). **(b)** RT-qPCR analysis of cardiac differentiation markers. Relative mRNA expression of *Tbx5* (first heart field marker), *Isl1* (second heart field/cardiac progenitor marker), *Tbx1* (pharyngeal mesoderm marker, negative control), and *Actc1* (cardiac structural marker) across days of differentiation, demonstrating sequential activation of the cardiac developmental program. **(c)** Representative flow cytometry analysis of differentiated hiPSC-CMs showing GFP expression versus cTnT staining. **(d)** Representative immunofluorescence image of an isolated bioincubated hiPSC-CM from rat heart demonstrating rod-shaped morphology with organized sarcomeric structures. Scale bar: 100 *µ*m. Green: GFP; red: cTnT. **(e)** Side-by-side comparison of bioincubated hiPSC-CM (left panels) and native rat CM (right panels). Scale bar: 50 *µ*m. Individual channels shown: GFP (green), cTnT (red), DAPI (blue, nuclear stain), and merged image. Both cell types display elongated morphology with organized sarcomeres and multinucleation. **(f)** Haralick correlation analysis of sarcomere organization in native rat cardiomyocytes (rCM, red) and bioincubated human iPSC-CMs (hCM, blue). Periodic correlation patterns indicate organized sarcomeric structures in both cell types. **(g)** Quantification of sarcomere length in native rat CMs (rCM) and bioincubated hiPSC-CMs (hCM). No significant difference was observed. Each point represents one cell; N = 5 animals. ns, not significant. **(h)** Relative standard deviation of sarcomere length within individual cells. Bioincubated hiPSC-CMs exhibited significantly higher intra-cellular variability compared to native rat CMs. \**p* < 0.05. **(i)** Cell circularity comparison. Lower values indicate more elongated, rod-shaped morphology. No significant difference was observed. ns, not significant. **(j)** Cell area comparison between rCM and hCM. No significant difference was observed. ns, not significant. **(k)** Gross image of explanted pig heart showing the intramyocardial injection site in the left ventricular free wall. **(l)** Histological sections of pig myocardium demonstrating engraftment of RFP-positive hiPSC-CMs (red signal) within the host myocardium. Engrafted cells display elongated morphology consistent with cardiomyocyte maturation. Scale bars: left, 200 *µ*m; right, 50 *µ*m.

RT-qPCR analysis demonstrated the expected temporal expression pattern of cardiac lineage genes, with early induction of cardiac progenitor markers (*Tbx5* and *Isl1*), followed by upregulation of structural cardiomyocyte markers such as *Actc1* (Fig. 1b). Flow cytometry demonstrated robust cTnT-positive cardiomyocyte differentiation (Fig. 1c).

To test the potential of neonatal hearts in CM maturation, GFP-labeled hiPSC-CMs were intramy-ocardially injected into neonatal rat hearts. After 4 weeks of bioincubation, hearts were harvested and enzymatically digested. COPAS large-particle sorting enabled recovery of live functional adult rod-shaped cardiomyocytes (up to 200 *µ*m in length) [6, 7]. Visual inspection confirmed that sorted cells retained intact morphology without obvious membrane damage (Fig. 1d). Immunofluorescence staining revealed structural sarcomere organization of mature adult cardiomyocytes compared to immature, less organized phenotypes of *in vitro* cultured hiPSC-CMs. Bioincubated hiPSC-CMs displayed elongated, rod-shaped morphology characteristic of adult cardiomyocytes (Fig. 1d). Staining for cTnT demonstrated well-organized sarcomeric structures with clear striation patterns (Fig. 1d, e). Bioincubated hiPSC-CMs exhibited binucleation, a hallmark feature of mature mammalian cardiomyocytes (Fig. 1e).

Sarcomere organization was evaluated using Haralick correlation analysis (Fig. 1f). Bioincubated hiPSC-CMs (hCM) demonstrated periodic correlation patterns consistent with organized sarcomeres, similar to adult rat cardiomyocytes (rCM). The sarcomere length measurements for both populations fell within the physiological range of 1.9–2.3 *µ*m reported for adult mammalian cardiomyocytes [14], with no significant difference between bioincubated hiPSC-CMs and native rat cardiomyocytes (Fig. 1g). Cell circularity showed no significant difference between groups, with both populations exhibiting values consistent with elongated rod-shaped morphology (Fig. 1i). Cell area was also comparable between the two populations (Fig. 1j). Despite similar average sarcomere dimensions, intra-cellular variability of sarcomere length remained significantly higher in bioincubated hiPSC-CMs compared to native rat cardiomyocytes (Fig. 1h; *p* < 0.05). This suggests that while bioincubated hiPSC-CMs achieve adult-range sarcomere length and cell geometry, the uniformity of sarcomere organization within individual cells has not yet reached the level of endogenous cardiomyocytes at 4 weeks of incubation.

To test feasibility of large-scale production of human adult CMs, we performed the bioincubation procedure in neonatal pigs by injecting RFP-labeled hiPSC-CMs into pig myocardium. Following bioincubation, hearts were analyzed by histological analysis. Gross inspection identified the injection site within the left ventricular myocardium (Fig. 1k). Histological sections demonstrated RFP-positive cells within the pig myocardium, confirming survival and engraftment of transplanted hiPSC-CMs (Fig. 1l). Engrafted cells displayed elongated morphology consistent with cardiomy-ocyte maturation. These preliminary findings provide proof-of-concept that hiPSC-CMs can survive and engraft in large animal hearts.

## DISCUSSION

In the present study, we show that bioincubation of hiPSC-CMs in neonatal hearts promotes maturation toward adult-like phenotypes. Bioincubated hiPSC-CMs isolated from rat hearts exhibited rod-shaped morphology, binucleation, and organized sarcomeric structures—features rarely observed in conventionally cultured hiPSC-CMs. Quantitative analysis revealed that bioincubated hiPSC-CMs achieve sarcomere length and cell circularity comparable to native rat cardiomyocytes. Furthermore, we provided preliminary evidence that hiPSC-CMs can successfully engraft in neonatal pig hearts, supporting feasibility of bioincubation in large animal models.

These findings are consistent with previous reports demonstrating hiPSC-CM maturation following *in vivo* transplantation: Kadota et al. reported hiPSC-CM structural maturation in rat hearts, with corresponding increases in sarcomere length and cell area [12]. Our study extends these findings by demonstrating maturation in neonatal rat hearts into an adult-like phenotype and by providing preliminary evidence in a large animal species. Although bioincubated hiPSC-CMs achieved comparable average sarcomere length to native cardiomyocytes, intra-cellular variability in sarcomere organization remained higher. Human cardiomyocytes mature more slowly than rodent cardiomyocytes even in the same *in vivo* environment [3, 12], and the higher variability we observed may reflect an intermediate maturation stage that could resolve with prolonged bioincubation. COPAS large-particle sorting offers a practical solution for isolating bioincubated hiPSC-CMs [6, 7].

Several limitations should be acknowledged. First, our characterization focused on morphological parameters; functional and transcriptomic assessments were not performed. Second, sample sizes were limited. Third, the pig data are preliminary, with histological confirmation of engraftment but without detailed morphometric analysis. Fourth, the absence of age-matched *in vitro* cultured hiPSC-CMs prevented quantitative attribution of maturation specifically to the bioincubation environment versus time-dependent maturation. Together, these results support bioincubation in neonatal hearts as a tractable *in vivo* platform for advancing hiPSC-CM maturation and justify future studies incorporating functional, transcriptomic, and long-term large-animal assessments.

## AUTHOR CONTRIBUTIONS

P.A. and C.K. conceived and designed the study. P.A. and N.H. performed the experiments. H.W. and P.A. analyzed the data. H.W., D.L., and C.K. wrote the manuscript. All authors reviewed and approved the final version.

## DISCLOSURE OF POTENTIAL CONFLICTS OF INTEREST

The authors declare no potential conflicts of interest.

## ACKNOWLEDGEMENTS

This research was supported by funding from NIH/NHLBI (R01HL156947, R01HL171205) and MSCRF (2017-MSCRFV-4049).

## Notes

### Competing Interest Statement

The authors have declared no competing interest.

